# NK cells orchestrate splenic cDC1 migration to potentiate antiviral protective CD8+ T cell responses

**DOI:** 10.1101/2020.04.23.057463

**Authors:** Sonia Ghilas, Marc Ambrosini, Jean-Charles Cancel, Marion Massé, Hugues Lelouard, Marc Dalod, Karine Crozat

## Abstract

A successful immune response relies on a tightly regulated delivery of the right signals to the right cells at the right time. Here we show that innate and innate-like lymphocytes use two mechanisms to orchestrate in time and space the functions of conventional type 1 dendritic cells (cDC1) in spleen. Early after murine cytomegalovirus infection, XCL1 production by lymphocytes with innate functions attracts red pulp cDC1 near IFN-γ-producing NK cells, generating superclusters around infected cells in the marginal zone. There, cDC1 and NK cells physically interact reinforcing their reciprocal activation. Targeted IL-12 delivery and IL-15/IL-15Rα transpresentation by cDC1 trigger NK cell activation and expansion. In return, activated NK cells deliver GM-CSF to cDC1, triggering their CCR7-dependent relocalization into the T cell zone. This NK cell-dependent licensing of cDC1 accelerates the priming of virus-specific CD8^+^ T cells. Our findings reveal a novel mechanism through which cDC1 bridge innate and adaptive immunity.

## Introduction

The success of an immune response against infections depends on timely coordinated actions of both the innate and adaptive immunities. In lymphoid organs, the delivery of the right signals to the right cells at the right time guarantees a rapid and efficient relay from innate cells to T cells, boosting specific and durable protective immunity. However, the mechanisms that come into play to orchestrate this tight cooperation are still poorly understood. Dendritic cells (DCs) are key in the activation of innate lymphoid cells (ILCs) and in the priming of T lymphocytes. How DCs are capable of delivering the help of the innate immune cells to naive T cells remain largely unknown.

Type 1 conventional DCs (cDC1) are critical in mounting cytotoxic CD8^+^ T lymphocyte (CTL) immune responses against extracellular pathogens (Merad et al., 2013) and cancers (Cancel et al., 2019; Wculek et al., 2020), in part due to their high efficacy at cross-presenting exogenous antigens to CD8^+^ T cells. Through their capacity to rapidly produce large amount of bioactive IL-12 (Dalod et al., 2002; Reis e Sousa et al., 1997), cDC1 are also good activators of innate lymphoid cells (ILCs), such as Natural Killer (NK) cells, and of T cell subsets with innate-like functions such as memory CD8^+^ T cells (Mashayekhi et al., 2011; Miyake et al., 2009; Alexandre et al., 2016). Regardless of their tissues of residency, cDC1 are phenotypically defined by their selective expression of the chemokine receptor XCR1 (Bachem et al., 2012; Crozat et al., 2011, 2010; Dorner et al., 2009). The unique known ligand for XCR1 in mice is XCL1, which is mostly expressed by NK cells and memory T cells at steady state, and further induced in these cell populations, and in CD8^+^ T and type 1 helper CD4^+^ T cells upon activation (http://www.immgen.org) (Crozat et al., 2010; Yamazaki et al., 2010). XCR1 expression on cDC1 potentiates CD8^+^ T cell priming and effector functions against infection and immunization (Crozat et al., 2010; Dorner et al., 2009), in part by orchestrating formation of clusters between cDC1 and XCL1-producing naive CD8^+^ T cells in lymphoid organs (Brewitz et al., 2017). XCR1 expression increases cDC1 infiltration in XCL1-expressing inflammatory tumors (Böttcher et al., 2018). XCR1 was also suggested to help cDC1 interpreting guiding cues provided by NK1.1^+^ or AsialoGM1^+^ cells (Böttcher et al., 2018), which include not only NK cells but also NK T cells and certain subsets of CD8^+^ and CD4^+^ T cells (Iigo et al., 1997; Trambley et al., 1999; Slifka et al., 2000; Yamanokuchi et al., 2005; Kosaka et al., 2007; Moore et al., 2008; Nour-Eldine et al., 2018). Although NK cells are a critical source of XCL1, the contribution of the XCL1/XCR1 axis in cDC1/NK cell interactions has not been rigorously determined. In addition, the mode of action of XCL1/XCR1 is still not completely understood: the XCL1/XCR1 axis could act by stabilizing cDC1/lymphocyte conjugates, or by promoting cDC1 microanatomical repositioning toward XCL1-producing cells.

cDC1 encompass the XCR1^+^CD8α^+^ cDC1 of lymphoid organs, and the peripheral XCR1^+^CD103^+^CD11b^-^ cDC1 residing in non-lymphoid tissues. Peripheral cDC1 convey tissue-derived antigens to naive T cells by continuously migrating through a CCR7-dependent mechanism, from the periphery to the tissue-draining lymph nodes (LNs) (Förster et al., 2012; Ohl et al., 2004; Saeki et al., 1999) to reach the deep T cell zone (Kitano et al., 2016). There, cDC1 make sequential physical contacts with CD4^+^ T cells and CD8^+^ T cells, facilitating the relay of the helper signal from the former subset to the latter, promoting protection against viruses (Hor et al., 2015; Eickhoff et al., 2015). In the spleen, the relationship between the different locations of cDC1 and the delivery of their functions is not as well understood as in the LN. The spleen is a highly compartmentalized lymphoid organ filtrating circulation for blood borne pathogens. The red pulp region is the first entry point into the spleen and mostly hosts cells with innate immune functions such as monocytes, macrophages, NK and NKT cells. The white pulp contains B cell follicles encircling T cell zones, where naïve T cells are confined. The bridging channel links the T cell zones to the red pulp, facilitating passage between the two compartments. Contrary to LNs where cDC1 remain restricted to the T cell zone, cDC1 in the spleen are found scattered in all compartments with the exception of B cell follicles (Alexandre et al., 2016; Calabro et al., 2016; Dorner et al., 2009). This regional segregation of cDC1 raises the question of the division of labor within the cDC1 population in the spleen: the red pulp cDC1 subset may be more prone to interact with innate cells, whereas the T cell zone cDC1 subset may be poised to initiate adaptive responses. Interestingly, systemic administration of poly(I:C), red blood cells, or *Listeria* triggers red pulp cDC1 repositioning in the marginal zone of the white pulp, and in the T cell zone where antigen-bearing mature cDC1 have more chance to interact with naïve T cells (Alexandre et al., 2016; Liu et al., 2019; Idoyaga et al., 2009; Calabro et al., 2016). CCR7 is suggested to be a master regulator of the migration of red pulp cDC1 into the T cell zone (Calabro et al., 2016). However, the critical time frame during which red pulp cDC1 acquire the guiding signaling to move into the T cell zone is still unknown, and the molecular mechanisms regulating their mobility inside the spleen remain obscure.

To address this, we used mouse cytomegalovirus (MCMV), because its infection induces a robust immune response encompassing the activation of both NK cells, and CD8^+^ T cells, which reside in the red pulp and the T cell zone respectively, and which activation strongly relies on cDC1 functions (Alexandre et al., 2014; Brizić et al., 2014). cDC1 contribute to activate NK cells through IL-12 (Dalod et al., 2002; Krug et al., 2004), a cytokine critical for NK cell IFN-γ production and host resistance (Nguyen et al., 2002; Pien et al., 2000). It has also been suggested that cDC1 regulated the expansion of antiviral Ly49H^+^ NK cells, through a mechanism proposed to involve IL-12 and IL-18 (Andrews et al., 2003). Moreover, in this model, the priming of antiviral CD8^+^ T cells occurs mainly through cross-presentation of viral antigens by cDC1 (Busche et al., 2013; Snyder et al., 2010; Torti et al., 2011). Finally, as early as one day after MCMV infection, before their full activation, NK cells produce the XCL1 chemokine (Dorner et al., 2004; Bezman et al., 2012). Therefore, MCMV allowed us to assess the specific contribution of the XCL1/XCR1 axis in cDC1/NK cell interactions *in situ*, a phenomenon that has never been visualized before. Thus, our advanced knowledge of the nature and kinetics of the activation of cDC1, NK cells and CD8^+^ T cells during MCMV infection made it a particularly well suited model to examine how cDC1 deliver their specific functions in time and space to NK and CD8^+^ T cells to orchestrate protective innate and adaptive immune defenses.

## Results

### XCR1 on cDC1 promotes NK cell activation, micro-anatomical repositioning and proliferation

To address the contribution of the XCL1/XCR1 axis in promoting cDC1/NK cell interactions, we first used the polyinosinic–polycytidylic acid (poly(I:C)), a synthetic analogue of double stranded (ds) RNA. Poly(I:C) administration induces a strong NK cell activation (Akazawa et al., 2007; McCartney et al., 2009), in part through direct contacts with DCs (Akazawa et al., 2007; Beuneu et al., 2009; Ebihara et al., 2010; Kasamatsu et al., 2014), and through stimulation by cDC1-derived IL-12 (McCartney et al., 2009; Miyake et al., 2009; Perrot et al., 2010). We found that depletion of cDC1 or of NK cell prior to poly(I:C) stimulation nearly abrogated NK cell IFN-γ production and cDC1 maturation respectively (Fig. S1B). Lack of *Xcr1* expression also decreased cDC1 and NK cell responses to poly(I:C) (Fig. S1C-E). These results showed the critical contribution of the XCL1/XCR1 axis to the cooperation between cDC1 and NK cells, promoting their reciprocal activation in the spleen in response to poly(I:C).

We next investigated whether a similar phenomenon occurs during productive viral infection by analyzing the impact of *Xcr1* genetic inactivation on NK cell responses to MCMV infection. At 40h post-infection, the absence of XCR1 altered NK cell activation and effector responses, as assessed by their CD69 expression (Fig. 1A), IFN-γ production (Fig. 1B) and upregulation of the cytotoxic molecule GzmB (Fig. 1C). At steady state, in the spleen, NK cells mostly reside in the red pulp (Fig. S1F) (Gregoire et al., 2008; Jaeger et al., 2012; Walzer et al., 2007). Upon MCMV, NK cells formed clusters in the marginal zone, especially around MCMV-infected cells, where they were activated as demonstrated by *in situ* IFN-γ staining (Fig. 1D). XCR1 expression on cDC1 increased the numbers of NK cell clusters at the edge of the marginal zones (Fig. 1E-F). Furthermore, XCR1 also promoted Ly49H^+^ NK cell proliferation as assessed by the upregulation of KI67 marker (Fig. 1G), and late expansion in the spleen (Fig. 1H). Altogether, these results showed that NK cell antiviral responses against MCMV strongly depended on the XCL1/XCR1 axis, and most likely on specific cDC1 functions.

**Figure 1:**
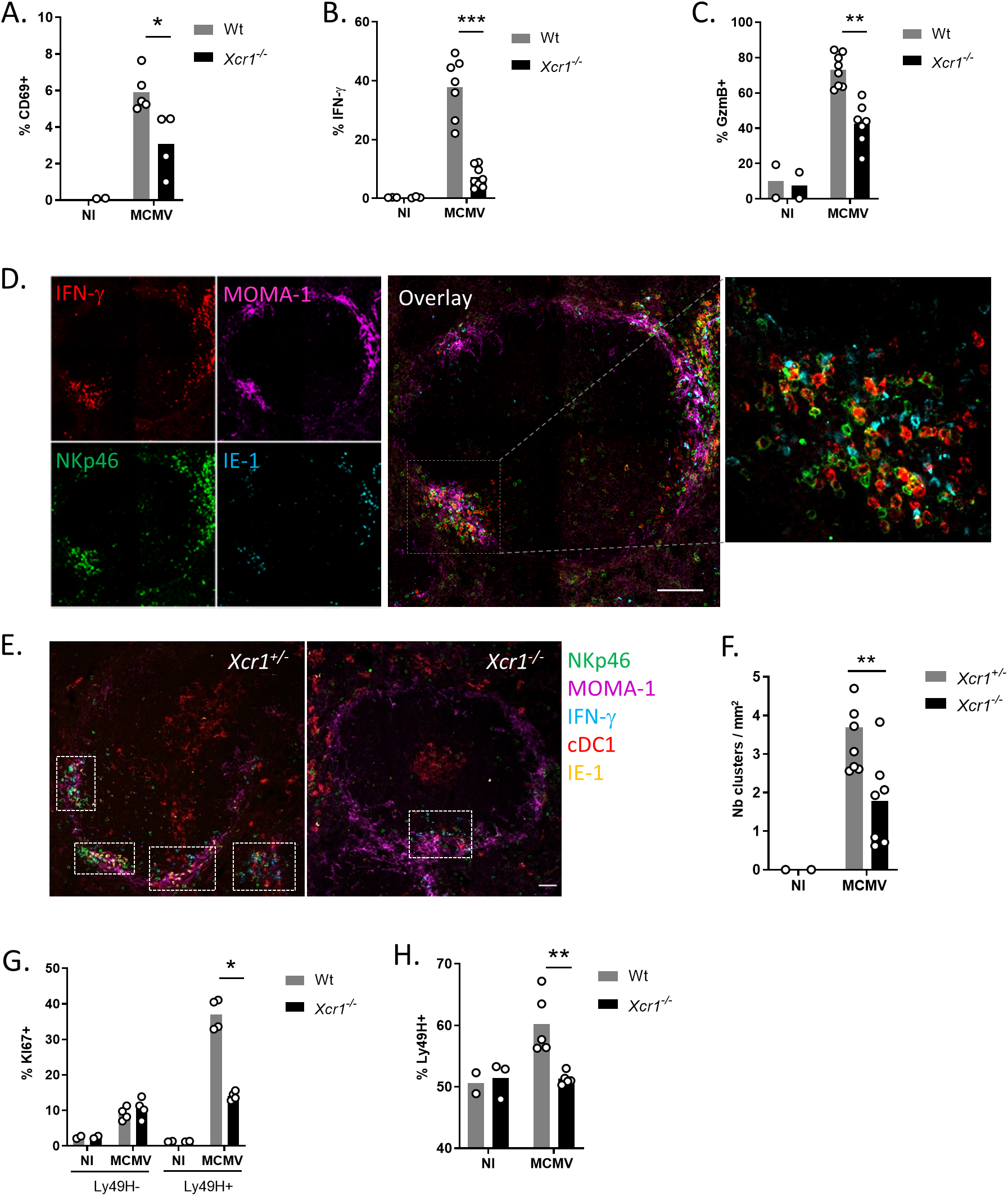
XCR1 promotes NK cell activation and redistribution in the spleen upon MCMV infection. Spleens of *Xcr1*^*-/-*^ mice and Wt controls were harvested 40h **(A-F)**, 4 days **(G)** and 5 days **(H)** post-MCMV infection. Splenic NK cells (NK1.1^+^TCRβ^-^) were stained for CD69 **(A)**, and intracellularly for IFN-γ **(B)**, and GzmB **(C)** directly *ex vivo*. **D)** Visualization of activated NK cell clusters in the spleens of MCMV-infected mice. Scale bar: 50μm. **E)** Analysis of NK cell clusters in *Xcr1*^-/-^ *vs* littermate controls upon MCMV. Spleen sections were prepared from *Karma*^*Cre*^;*Rosa26*^*tdRFP*^;*Xcr1*^*-/-*^ mice and *Karma*^*Cre*^;*Rosa26*^*tdRFP*^;*Xcr1*^*+/-*^ controls and stained for NKp46 (green), MOMA-1/CD169 (purple), IFN-γ (cyan), tdRFP (red) and IE-1 (yellow). Inserts show examples of clusters areas as defined in the material and methods. NK cell clusters were identified as groups of at least 10 IFN-γ^+^ cells, encompassing NKp46^+^ cells and gathered around MCMV-infected cells (IE-1^+^) in the marginal zone (MOMA-1^+^). Scale bar: 50μm. **F)** Quantification of NK cell clusters in the spleens of MCMV-infected *Karma*^*Cre*^;*Rosa26*^*tdRFP*^;*Xcr1*^*-/-*^ mice and their littermate *Xcr1*^*+/-*^ control 40h post-infection. **G)** Splenic NK cells (NK1.1^+^TCRβ^-^) were stained intracellularly for KI67 4 days post-infection. **H)** Proportion of Ly49H^+^ NK cells in Wt vs *Xcr1*^-/-^ mice 5 days post infection. One experiment representative of at least 4 independent ones with at least 4 mice per infected group is shown, except for B-E, where two experiments with at least 3 mice per group were pooled. NI, non-infected; *, p<0.05; **, p<0.01; ***, p<0.001.

### XCR1 participates in cDC1 redistribution toward NK cells, promoting their production of IL-12

We sought to determine to what extend XCR1 regulated cDC1 responses to MCMV infection, in particular their capacity to produce IL-12 and to mature phenotypically. As compared to Wt controls, at 40 h post-infection, XCR1-deficient mice showed a strong decrease in the proportion of cDC1 producing IL-12 (Fig. 2A), likely explaining their decreased proportion of IFN-γ^+^ NK cells (Fig. 1B) since cDC1 were the overwhelming source of IL-12 production amongst DC subsets (Fig. 2A and Fig. S2A). Interestingly, upregulation of the maturation markers CD86, CD80 and CD40 on cDC1 was not significantly altered in XCR1-deficient mice 48h post-infection (Fig. 2B). At steady state, cDC1 are scattered in the red pulp, and concentrated in T cell area in the spleen (Alexandre et al., 2016; Yamazaki et al., 2013) (Fig. S2B). We wondered whether MCMV infection induced the relocalization of splenic cDC1 to the NK cell islets at the sites of viral replication in the marginal zone, and whether this relied on XCR1. To track cDC1 *in situ*, we bred XCR1-deficient animals with *Karma*^*Cre*^;*Rosa26*^*tdRFP*^ mice in which the expression of the tandem dimer red fluorescent protein (tdRFP) selectively traced splenic CD8α^+^ cDC1 (Mattiuz et al., 2018), and we compared *Xcr1*^*-/-*^ animals to *Xcr1*^*+/-*^ littermate controls. In control mice, several cDC1 were observed inside each NK cell cluster, where they harbored a dendritic morphology (Fig. 2C). In the absence of XCR1, cDC1 were less abundant in clusters of activated NK cells, and were mostly round shaped (Fig. 2C). This differential repositioning of cDC1 between *Xcr1*^*-/-*^ vs littermate control mice was confirmed by an unbiased quantification of tdRFP intensity in individual NK cell clusters (Fig. 2D), whereas the mean tdRFP intensity on the whole spleen sections remained the same between both groups (Fig. S2C). To determine *in situ* whether those cDC1 that are in contact with NK cells were a source of IL-12, we bred the *Karma*^*Cre*^;*Rosa26*^*tdRFP*^ mice with *Il12eYFP*^*+*^ animals reporting IL-12-p40-expressing cells as fluorescent for eYFP (Reinhardt et al., 2006). In the marginal zone clusters, IL-12^+^ cDC1 were making contacts with NK cells (Fig. 2E), many of which were polarizing their IFN-γ at the interface with cDC1 (Fig. 2F and Video S1). This observation confirmed the formation of stimulatory synapses between DCs and NK cells that had been previously described *in vitro* (Borg et al., Blood 2004). Moreover, when IL-15Rα expression was selectively inactivated in cDC1 to abrogate their IL-15 trans-presentation capacity, there was a significant reduction of NK cell number in the spleen 4 days post-infection (Fig. 2G). Altogether, these results showed that the XCL1/XCR1 axis reinforced the attraction of cDC1 toward NK cell clusters, thereby promoting a feedforward loop enhancing physical interactions between these two cell types, and a reciprocal activation, at least in part through local delivery of IL-12 and of IL-15/IL-15Rα by cDC1 to NK cells.

**Figure 2:**
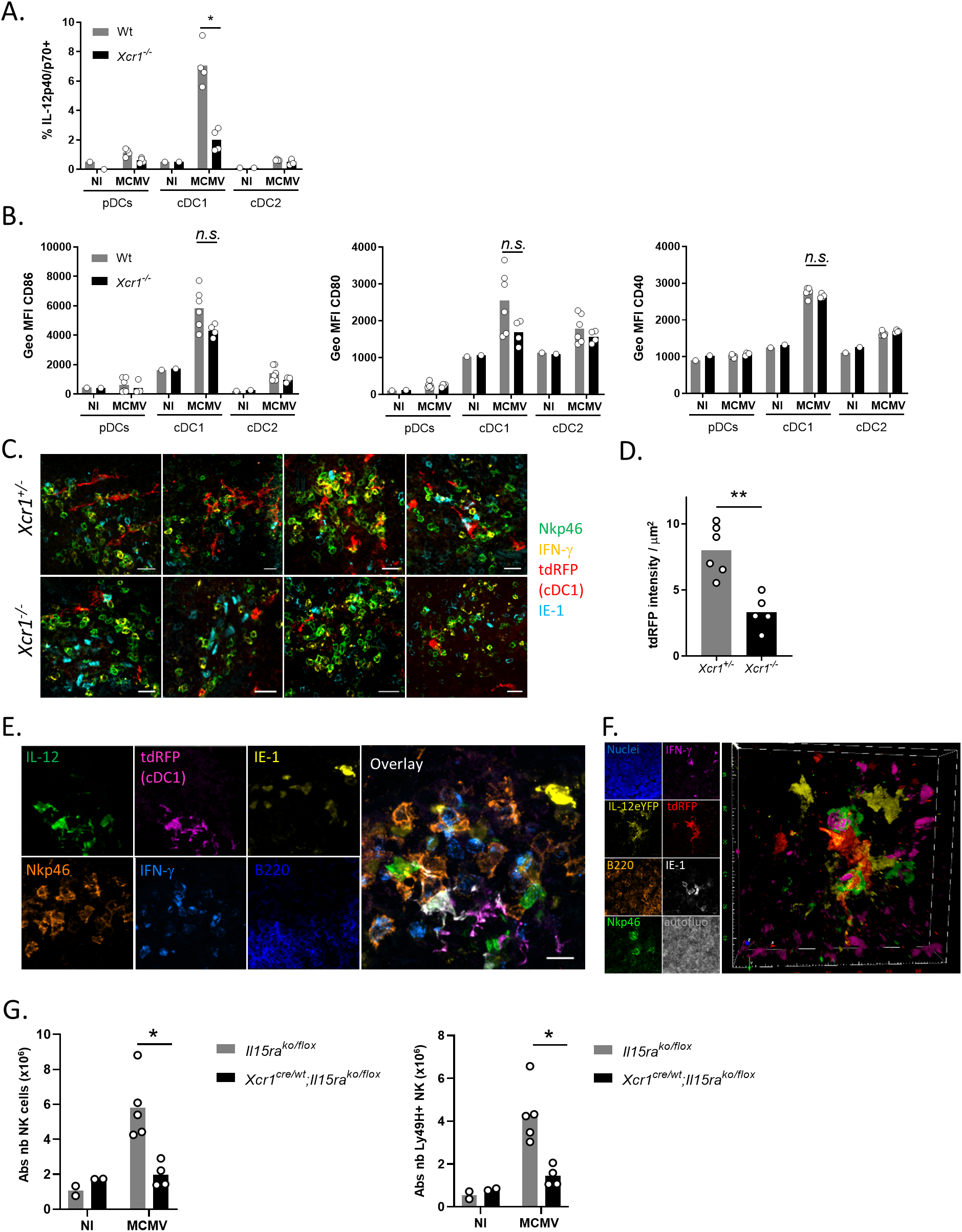
XCR1 regulates IL-12 production by cDC1, and their positioning into clusters where IL-12-producing cDC1 contact NK cells. **A)** Spleens were harvested at 40h **(A; C-F)** and 48 hours **(B)** post MCMV infection. **A-B)** Analysis of IL-12p40/p70 production **(A)** and CD86, CD80 and CD40 expression **(B)** by splenic DCs of *Xcr1*^*-/-*^ and Wt animals. One experiment representative of at least 4 independent ones with at least 4 mice per infected group is shown. NI, non-infected; *n*.*s*., non-significant; *, p<0.05. **C-D)** Visualization (**C**) and quantification (**D**) of cDC1 presence in NK cell clusters in *Karma*^*Cre*^;*Rosa26*^*tdRFP*^;*Xcr1*^*-/-*^ mice and their *Xcr1*^*+/-*^ littermate controls 40h post-infection. Spleen sections (scale bar, 20μm) were stained for NKp46 (green), IFN-γ (cyan), tdRFP (red) and IE-1 (yellow) and individual clusters were isolated as shown in **(C)**, before tdRFP intensity per μm^2^ was computationally quantified in each cluster **(D)**. Two independent experiments with 2-3 mice per infected group were pooled. **, p<0.01. **E)** Analysis of IL-12-producing cDC1 in NK cell clusters in *Karma*^*Cre*^;*Rosa26*^*tdRFP*^;*Il12eYFP*^*+*^ mice 40h post MCMV infection. The color obtained from overlaying green IL-12-eYFP^+^ on purple tdRFP-expressing cDC1 is white. Scale bar, 20μm. **F)** IL-12-eYFP-expressing cDC1 make physical contacts with IFN-γ-producing NK cells in marginal ring clusters. The micrographs shown **(E-F)** are representative of the analyses of 6 mice from two independent experiments. **G)** Analysis of NK cell absolute number 4 days post-infection in spleen of *Xcr1*^*Cre/wt*^;*Il15ra*^*KO/flox*^ mice and their respective controls (*Il15ra*^*KO/flox*^). One experiment representative of at least 2 independent ones with at least 4 mice per infected group is shown. NI, non-infected; *n*.*s*., non-significant; *, p<0.05.

### XCL1 from NK cells is not necessary to promote cDC1/NK cell contact interactions

In order to confirm that NK cells were a major cellular source of XCL1 upon MCMV infection, we examined *Xcl1* expression in total spleen of infected animals. Because commercially available anti-XCL1 antibodies did not reliably stain XCL1 *ex vivo* in our hands, we generated an *Xcl1-mTfp1*^*fl/fl*^ mouse model by knocking a LoxP-exon3-IRES-mTfp1-LoxP cassette in frame in the *Xcl1* gene (Fig. S3A). This construct was made such that 1) expression of the monomeric Teal fluorescent protein mTFP1 allowed the genetic tracing of all *Xcl1*-expressing cells; 2) the recombination of LoxP sequences by the CRE recombinase inactivated both *Xcl1* and *mTfp1* expression selectively in Cre^+^ cells. At steady state, mTFP1 expression was the strongest in ILC1, NK cells and NKT cells (Fig. 3A), with NK cells being the major cell subset expressing *Xcl1* (49.9% of all mTFP1^+^ splenocytes), followed by CD44^+^CD8^+^ T cells (13.1%) and CD44^+^CD4^+^ T cells (12.4%), which encompassed essentially memory αβ T cells (Fig. 3C). Although NK cells remained the major source of *Xcl1* upon MCMV infection (44.3%) (Fig. 3B and 3D), the inactivation of *Xcl1* selectively in NK cells and partly in ILC1 (Fig. S3B) did not decrease their activation including their capacity to produce IFN-γ, contrary to what happened in *Xcr1*^-/-^ animals (Fig. 3E). These results implied that memory/activated T cells, NKT cells and/or eventually ILC1 can compensate for the lack of XCL1 production by NK cells during MCMV infection, with all these XCL1-producing cells likely acting together to attract cDC1 in specific foci around infected cells in the marginal zone of the spleen.

**Figure 3:**
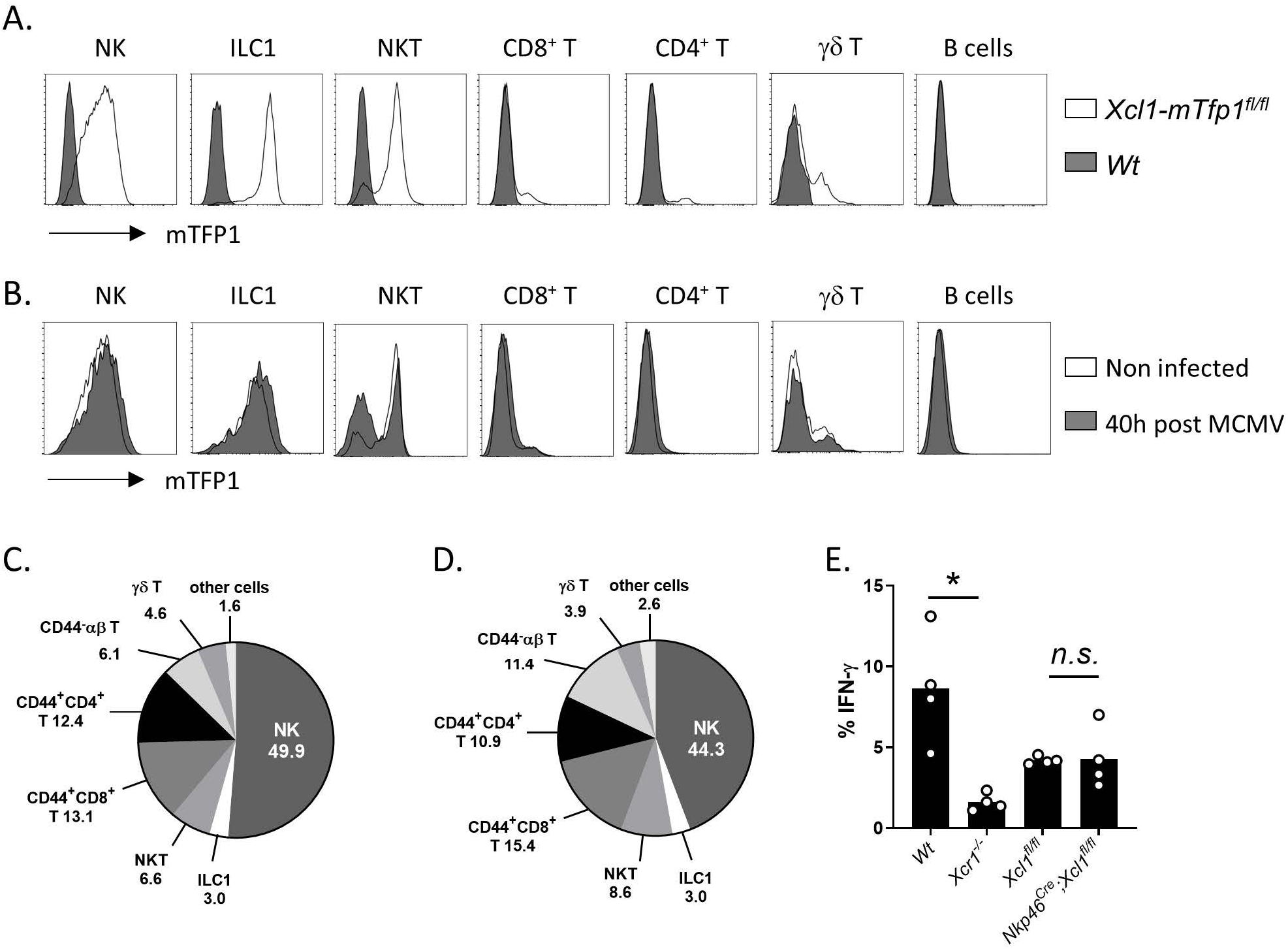
NK cells are a main source of XCL1 but not a critical one upon MCMV infection. **A-B)** Analysis of mTFP1 expression in splenocytes of Wt (dark grey) and *Xcl1-mTfp1*^*fl/fl*^ (white) mice at steady state **(A)** or 40h post-infection **(B)**. Cells were gated as follow: NK cells (TCRβ^-^ CD19^-^ NK1.1^+^), ILC1 (NK1.1^+^ TCRβ^-^ CD19^-^ CD127^+^), NKT cells (CD19^-^ CD1d^+^), CD8^+^ T (CD1d^-^ NK1.1^-^ TCRβ^+^ CD19^-^ CD8^+^), CD4^+^ T (CD1d^-^ NK1.1^-^ TCRβ^+^ CD19^-^ CD4^+^), γδ T cells (CD1d^-^ NK1.1^-^ CD3ε^+^ TCRβ^-^ TCRγδ^+^) and B cells (NK1.1^-^ TCRβ^-^ CD19^+^). One representative experiment of 4 independent ones with at least three mice per group is shown. **C-D)** Proportion of cell populations within mTFP1 positive cells at steady state **(C)** and 40h post-MCMV infection **(D)**. Others: sum of all the other cell subsets not detailed in the pie charts. One representative experiment of 4 independent ones with at least three mice per group is shown. **E)** IFN-γ production by NK cells in *NKp46*^*Cre*^;*Xcl1*^*fl/fl*^, *Xcl1*^*fl/fl*^, Wt and *Xcr1*^-/-^ mice 40h post-infection. One representative experiment of two independent ones with at least four mice per group is shown. n.s., non-significant; *, p<0.05.

### cDC1/NK cell interaction promote cDC1 redistribution from the red pulp to the T cell zone via the bridging channel

We set up a quantitative analysis to investigate the migratory behavior of cDC1 during the first 48 hours following MCMV infection. We used *Karma*^*Cre*^;*Rosa26*^*tdRFP*^ mice that we stained with MOMA-1 and B220 to delineate marginal zones, T cell zone and the bridging channels on whole spleen tissue sections (Fig. S4), and we quantified tdRFP intensity as a readout of cDC1 presence in each of these compartments. In *Xcr1*^*+/-*^ mice, cDC1 redistribution in the marginal zone peaked at 40h post-infection, and then cDC1 moved in the bridging channel at 44h to reach the T cell zone at 48h (Figure 4A). XCR1 inactivation compromised this migratory pattern of cDC1, which at 48h relocalized in the marginal zone rather than in the T cell area (Fig. 4A). We further investigated the molecular mechanisms providing guidance cues for marginal cDC1 to enter the T cell zone. The Epstein-Barr virus–induced gene 2 (EBI2; also known as GPR183) is a Gαi-coupled chemoattractant receptor required for cDC2 positioning in the bridging channel and T cell zone in response to 7α,25-(7α,25-HC) and 7α,27-(7α,27-HC) hydroxycholesterols (Lu et al., 2017; Yi and Cyster, 2013). *Gpr183* expression was not upregulated in *Xcr1*^*-/-*^ cDC1 upon MCMV infection (Fig. 4B), and, as a consequence, migration of cDC1 isolated from MCMV-infected *Xcr1*-deficient animals was completely abrogated in response to 7α,25-HC (Fig. 4C). These data supported the decreased cDC1 relocalization in *Xcr1*-deficient mice in the bridging channel at 44h post-infection (Fig. 4A). The chemokine receptor CCR7 also partly orchestrates DC movement into the T cell area in the spleen (Calabro et al., 2016; Gunn et al., 1999; Umemoto et al., 2012; Yi and Cyster, 2013). CCR7 cognate ligands, CCL19 and CCL21, are expressed on fibroblast reticular cells (FRC) in the bridging channel and T cell zone (Luther et al., 2000; Bajénoff et al., 2008). We found that *Xcr1*-deficient cDC1 did not express CCR7 protein at levels equivalent to littermate controls at 48h post-infection (Fig. 4D), which was the time when cDC1 relocalization peaked in the T cell zone in control mice (Fig. 4A). These results show that the XCR1-dependent repositioning of cDC1 within NK cell clusters in the marginal zone early after MCMV infection regulate their expression of molecules, such as Gpr183 or CCR7, that are key for cDC1 to interpret environmental guiding cues present across the bridging channel.

**Figure 4:**
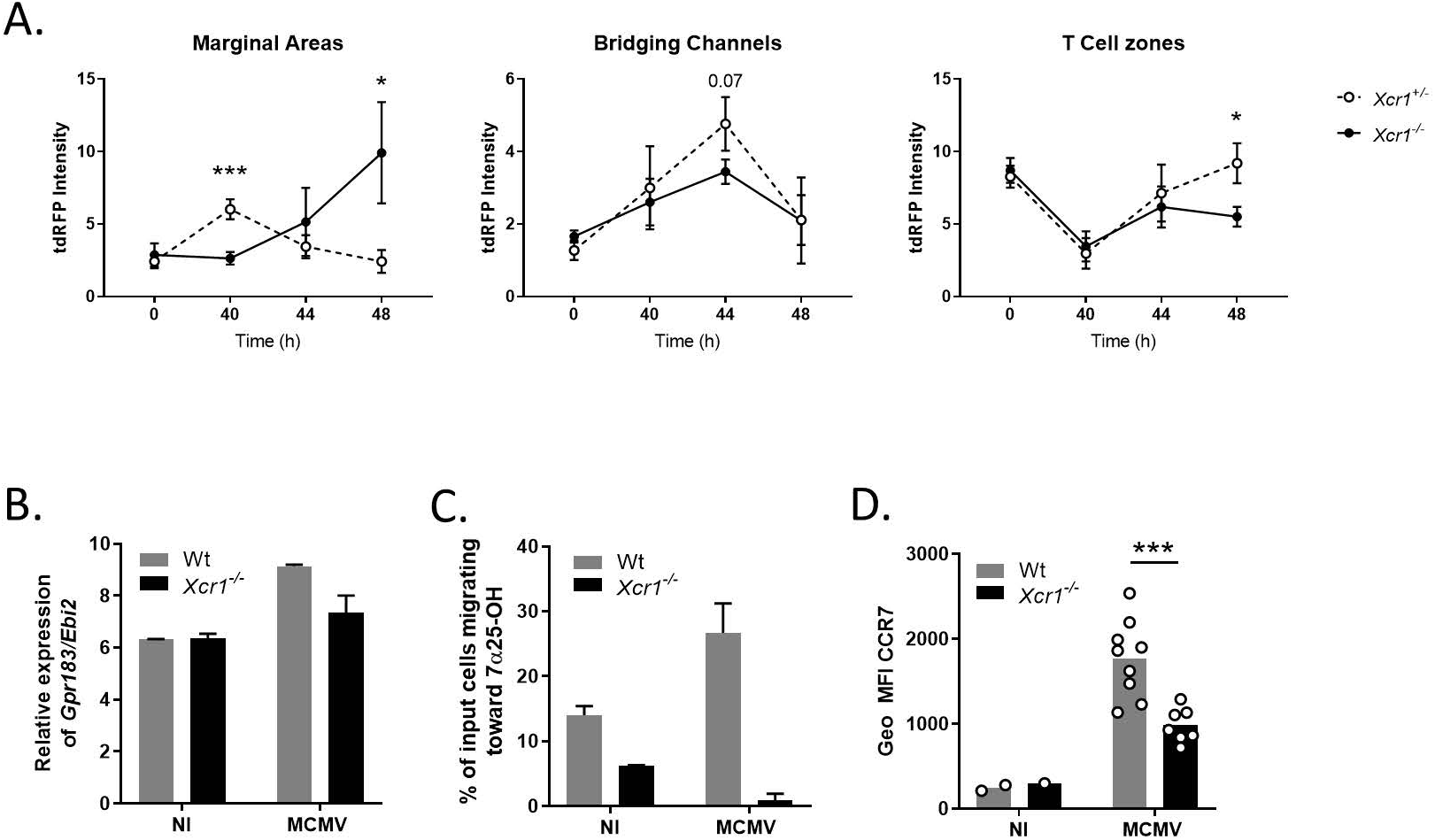
Upon MCMV infection, the spatiotemporal relocalization of cDC1 in the different regions of the spleen is associated with the regulation of EBI2 and CCR7 expression. **A)** Quantification of tdRFP intensity as a read out of cDC1 presence in marginal areas, bridging channels and T cell zones of *Karma*^*Cre*^;*Rosa26*^*tdRFP*^;*Xcr1*^*-/-*^ and their *Xcr1*^*+/-*^ littermate control, during the course of MCMV infection. MOMA-1^+^ ring defined marginal areas, B220 staining delineated T cell areas, and breaks in both B220 and MOMA-1 stainings defined bridging channels (see Fig. S4). Each dot (+/-SD) represents a pool of two experiments with 2-3 mice per group. Statistical test: 2-way ANOVA; *, p<0.05; ***, p<0.001. **B)** Relative expression of the *Gpr183*/*Ebi2* gene from transcriptomic analysis of splenic cDC1 sorted from Wt vs *Xcr1*^-/-^ 38h post-infection. NI, Non-infected. **C)** Transwell migration assay of cDC1 in response to the EBI2 ligand, 7α25OH oxysterol. DCs from Wt and *Xcr1*^-/-^ spleens were enriched by optiprep gradient 40h post-infection. NI, non-infected. **D)** Analysis of CCR7 expression on cDC1 in Wt and *Xcr1*^-/-^ mice 48h post-infection. Two independent experiments with at least 3 mice per infected group were pooled. NI, non-infected. *, p<0.05; ***, p<0.001.

### CCR7 upregulation on cDC1 depends on physical contacts with activated NK cells

Because XCR1-dependent cDC1 relocalization into NK cell clusters in the marginal zone was the major event preceding cDC1 migration into the T cell zone, we hypothesized that this step in the antiviral immune response might allow cDC1 to receive specific signals critical for their CCR7 upregulation. Indeed, NK cells isolated from MCMV-infected animals efficiently triggered CCR7 expression on immature cDC1 in an *in vitro* co-culture experiment, without changing their phenotypic maturation as assessed by CD86 expression (Fig. 5A). This NK cell-driven upregulation of CCR7 on cDC1 was significantly reduced when these two cell types were physically separated by a transwell (Fig. 5B), indicating that cDC1 need to closely contact activated NK cells to induce their expression of CCR7. On spleen sections, in NK cell clusters, the CCR7 antibody mostly stained cDC1 that touched IFN-γ-producing NK cells (Fig. 5C), confirming the hypothesis that physical contacts between cDC1 and NK cells were required for the former to acquire the capacity to migrate to the T cell area.

**Figure 5:**
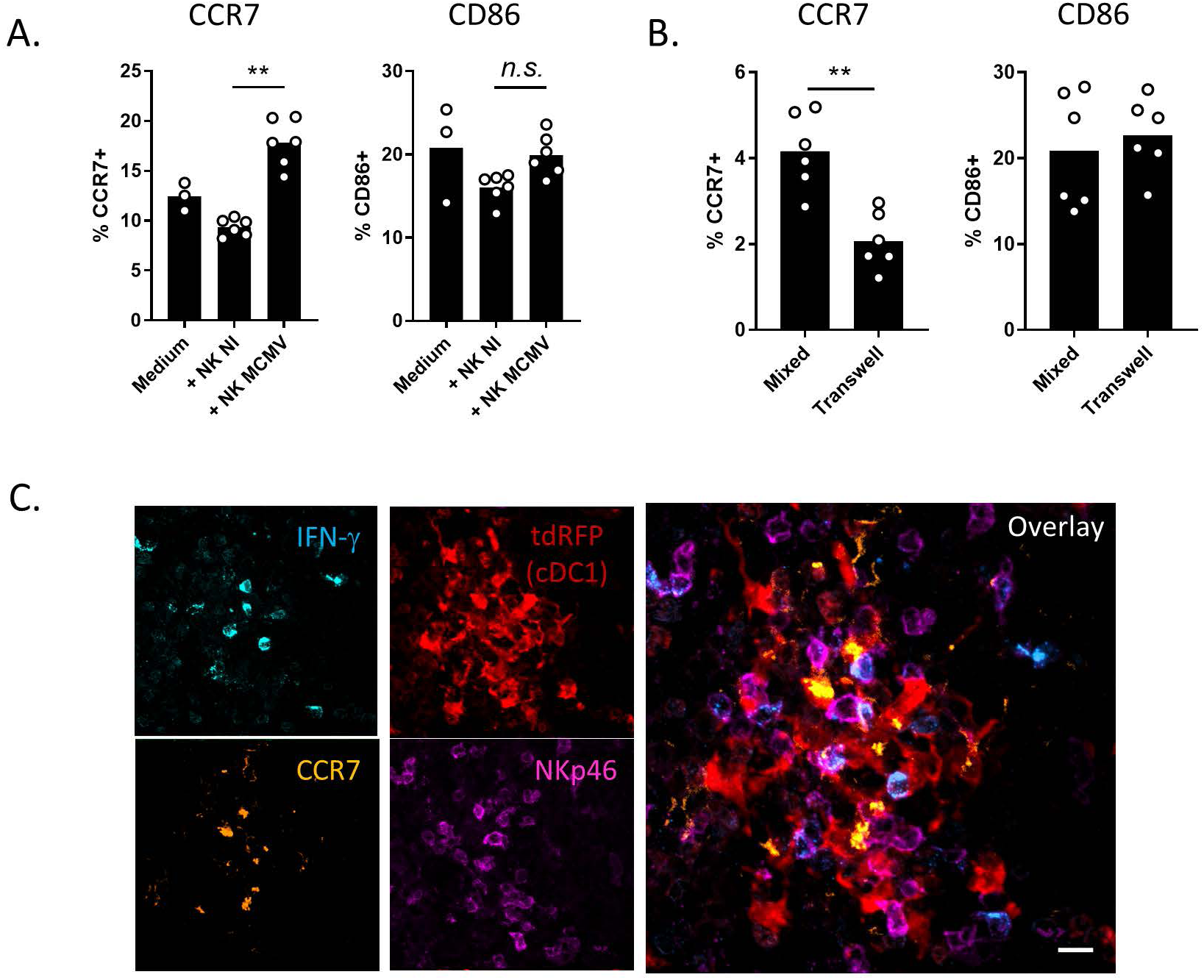
Activated NK cells upregulate CCR7 on cDC1 in a cell/cell contact dependent manner. **A-B)** Analysis of CCR7 and CD86 expression on cDC1 when in co-culture for 7h with NK cells from spleens of non-infected Wt mice or from mice infected for 40h with MCMV. cDC1 and NK cells were co-cultured either in the same well **(A)** or separated by a 0.4um diameter pores bearing transwell apparatus **(B)**. One representative experiment of two independent ones with at least four mice per group is shown. *n*.*s*., non-significant; *, p<0.05. Analysis of CCR7 expression on cDC1 in NK cell clusters 40h post MCMV infection. Spleen sections of *Karma*^*Cre*^;*Rosa26*^*tdRFP*^ mice were stained for IFN-γ, NKp46, CCR7 and tdRFP (cDC1). This cluster was imaged in the marginal area. The micrograph shown is representative of the analyses of 4 mice, from two independent experiments. Scale bar, 10μm.

### GM-CSF contributes to the upregulation of CCR7 expression on cDC1

We next sought for the signals provided by activated NK cells and promoting cDC1 upregulation of CCR7. *In vivo* neutralization of IFN-γ starting prior to MCMV infection did not decrease CCR7 upregulation on cDC1 (Fig. S5A). CCR7 upregulation on cDC1 48h after MCMV infection was correlated with a decrease in XCR1 expression (Fig. S5B), but supplementation of FLT3-L differentiated bone marrow-derived DCs with recombinant XCL1 did not lead to CCR7 upregulation the *in vitro* derived cells equivalent to splenic cDC1 (eqcDC1) (Fig. S5C). We then focused on the granulocyte-macrophage colony-stimulating factor (GM-CSF, encoded by the *Csf2* gene). GM-CSF was reported to improve eqcDC1 survival and functions in FLT3-L bone marrow cultures (Sathe et al., 2011; Zhan et al., 2011). The stimulation of eqcDC1 with a low concentration of GM-CSF was sufficient to induce a significant upregulation of their CCR7 expression (Fig. S5D). Interestingly, during MCMV infection, *Csf2* transcription followed a kinetic similar to that of *Ifng* and *Xcl1* in the spleen, reaching a peak at 40 h (Fig. 6A), corresponding to the time when cDC1 encountered NK cells in the marginal zone. At that time, among splenic lymphocytes, GM-CSF was mainly produced by NK cells (Fig. 6B). IL-12 stimulation efficiently triggered *Csf2* transcription in purified NK cells, when compared to other stimuli such as NK cell receptor engagement or soluble IL-15 (Fig. 6C). This suggested that, in the marginal zone cell clusters induced by MCMV infection, cDC1-derived IL-12 might be one of the main signals driving GM-CSF production by NK cells. We analyzed whether GM-CSF signaling on cDC1 was involved in CCR7 expression. Like splenic cDC1 (Fig. 5A), *in vitro* derived eqcDC1 upregulated CCR7 only when cultured with NK cells isolated from MCMV-infected mice (Fig. 6D). In this context, GM-CSFR-deficient (*Csf2rb*^*-/-*^) eqcDC1 showed a reduced capacity to upregulate CCR7 compared to Wt eqcDC1 (Fig. 6D). Moreover, the analysis of Wt:*Csfr2b*^*-/-*^ mixed bone marrow chimera mice infected by MCMV showed a specific defect of *Csfr2b*^*-/-*^ cDC1 to upregulate CCR7 (Fig. 6E). All these results strongly supported a critical contribution of NK cell-derived GM-CSF in promoting cDC1 migration from the marginal zone cell clusters to the T cell area within the spleen during MCMV infection, via CCR7 upregulation.

**Figure 6:**
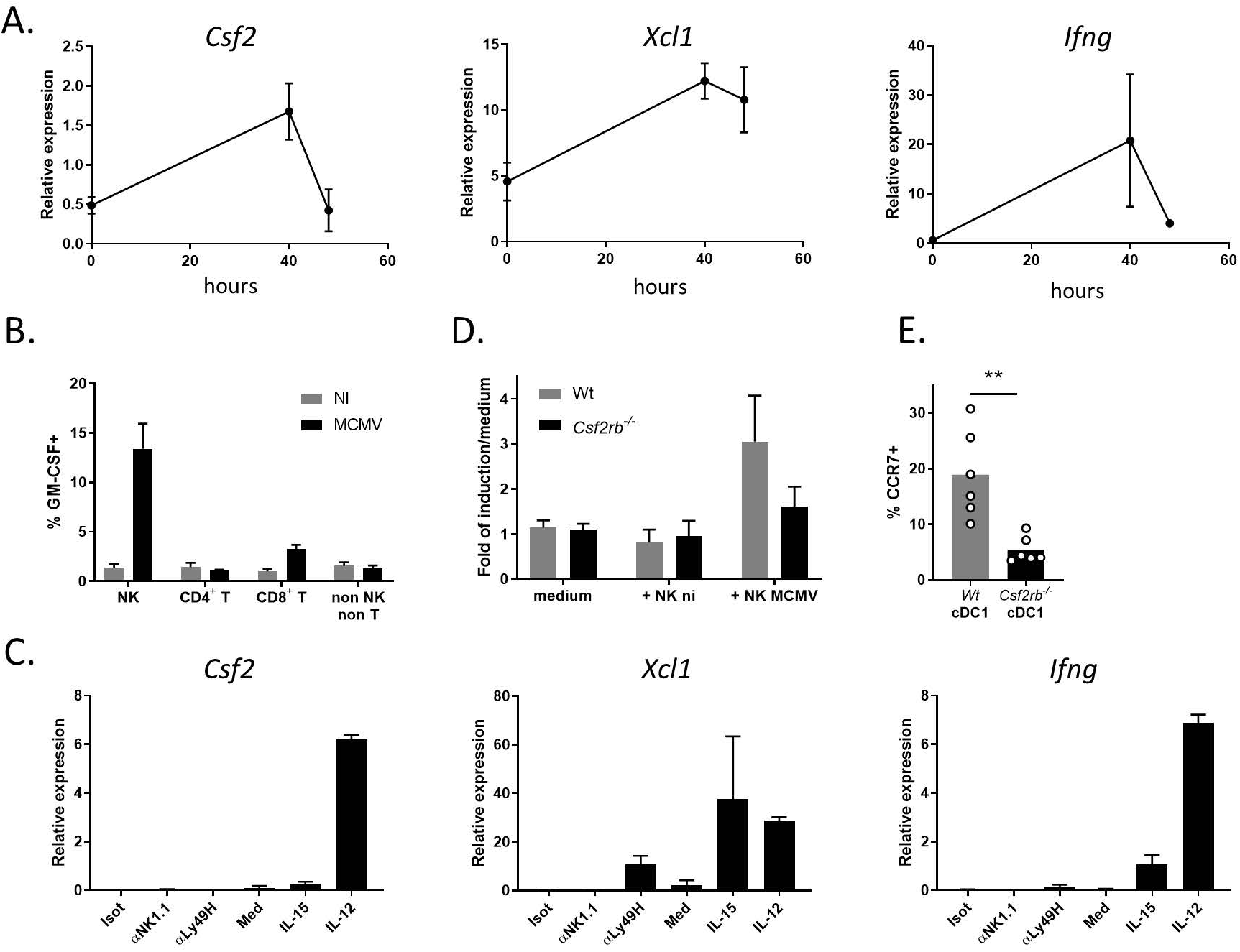
GM-CSF, largely produced by activated NK cells, is critical for CCR7 upregulation on cDC1. **A)** Kinetics of induction of *Csf2* (*Gmcsf*) expression in total spleen in the course of MCMV infection. Total RNA was extracted from spleens from Wt mice at the indicated times post MCMV infection, and RT-qPCR were performed using *Hprt* as housekeeping gene. One representative experiment of two independent ones with at least 4 mice per group is shown. **B)** Analysis of GM-CSF production by different lymphocyte populations 40h post MCMV infection. Splenocytes were intracellularly stained for GM-CSF. **C)** Analysis of *Csf2, Xcl1* and *Ifng* expression in purified NK cells activated *in vitro* with different stimuli. Analysis of CCR7 upregulation on *Csfr2*-deficient FLT3-L bone marrow derived cDC1 by activated NK cells. Wt and *Csfr2b*-deficient immature cDC1 were enriched from FLT3-L bone marrow cultures and co-cultures for 7h with NK cells purified from spleens of mice infected or not for 40h with MCMV. cDC1 were gated as CD11c^+^SiglecH^-^SIRPα^-^CD24^+^ cells. **E)** Analysis of CCR7 expression on Wt vs *Csf2rb*^*-/-*^ cDC1 from Wt:*Csf2rb*^*-/-*^ mouse chimeras 48h post infection. **, p<0.01. Error bars represent standard deviations.

### NK cell-mediated cDC1 relocalization to the T cell area promotes MCMV-specific CD8^+^ T cell responses and control of MCMV infection

We next wondered whether the decreased cDC1 migration to the T cell area of the spleen in *Xcr1*^*-/-*^ infected mice affected the induction of antiviral adaptive immunity. The expansion of CD8^+^ T cells specific for the non-inflationary viral peptide m45 was significantly reduced 6 days post-infection in absence of XCR1 (Fig. 7A). This observation confirmed the critical role of cDC1 in the cross-priming of antiviral CD8^+^ T cells during an acute MCMV infection (Fig. S6) (Busche et al., 2013; Snyder et al., 2010; Torti et al., 2011). Since the absence of XCR1 hampered both NK cell effector functions and expansion, and the induction of antiviral CD8^+^ T cell responses, we wondered whether this could have compromised viral control. We measured splenic and liver viral titers at 5 days post-infection, *i*.*e*. when they have normally become undetectable in Wt mice. Productive viral particles could still be detected in *Xcr1*-deficient animal (Fig. 7B). Thus, altogether, this study shows that, during MCMV infection, the XCL1/XCR1 axis promoted a crosstalk between cDC1 and NK cells leading to their mutual activation and to faster/higher downstream antiviral CD8^+^ T cell responses, overall strongly enhancing host resistance. Moreover, our study identified a new mechanism used by activated NK cells to boost adaptive immune responses to MCMV, a phenomenon described earlier by comparing the kinetic and intensity of antiviral CD8^+^ T cell responses between mice harboring or not NK cells able to specifically recognize and kill infected cells (Robbins et al., 2007).

**Figure 7:**
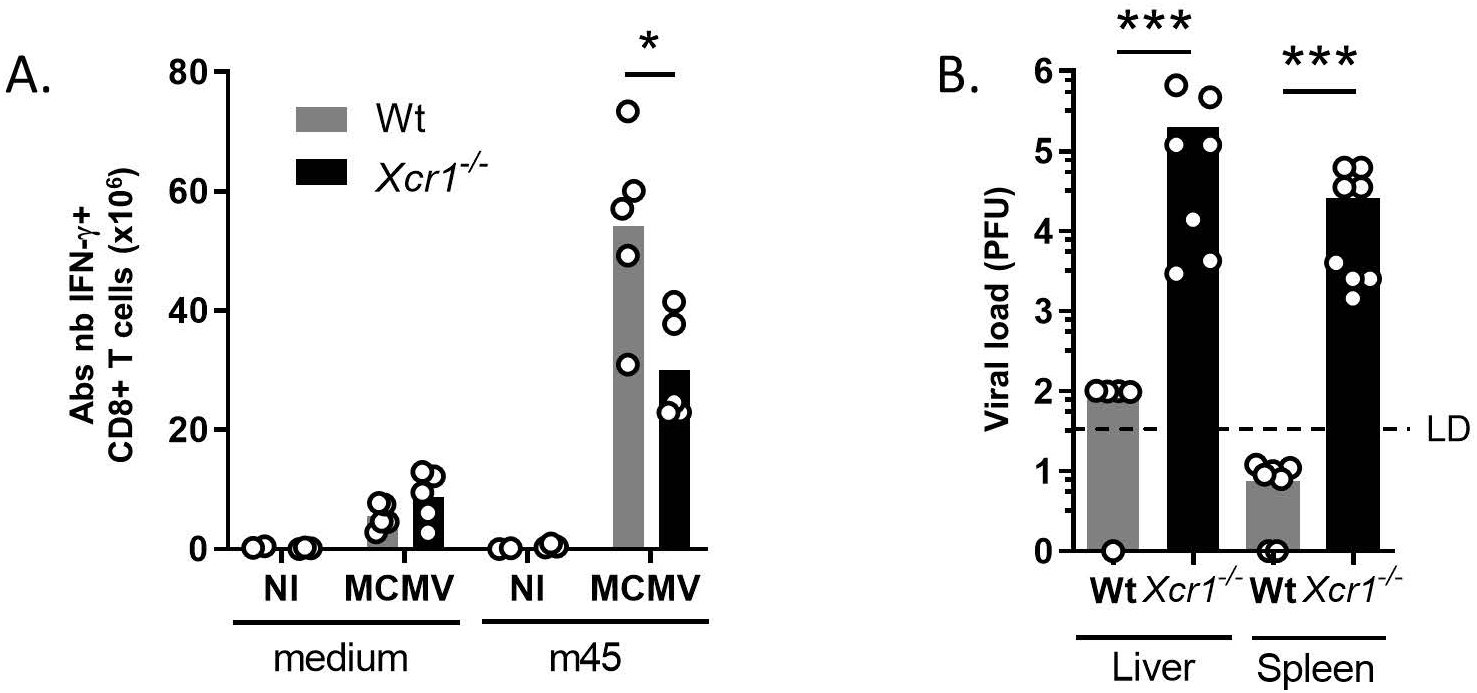
cDC1 relocalization defect compromise MCMV-specific CD8+ T cell responses and host resistance to MCMV infection. **A)** Analysis of m45-specific CD8^+^ T cell response in *Xcr1*^-/-^ and littermate controls 5 days post-MCMV infection. One representative experiment of two independent ones with at least 4 mice per group is shown. *, p<0.05. **B)** MCMV titers in spleens and livers of Wt and *Xcr1*^*-/-*^ mice 5 days post-MCMV infection (10^4^ PFU). Two independent experiments with at least 3 mice per infected group were pooled. NI, non-infected. ***, p<0.001.

## Discussion

Here we use MCMV infection to describe the molecular mechanisms conferring to NK cells the capacity to license cDC1 migration in the spleen, fast-forwarding CD8^+^ T cell priming and leading to efficient antiviral responses. The XCL1/XCR1 axis constitutes the foremost step that triggers the redistribution of red pulp cDC1 near activated NK cells and MCMV-infected cells in the marginal zone. The failure of cDC1 to reposition themselves in contact with NK cells curtails the antiviral defenses by limiting not only NK cell IFN-γ production and expansion, but also antiviral CD8^+^ T cell responses. We also show that tight contacts between cDC1 and NK cells in the marginal zone are necessary to set up a productive crosstalk with the delivery of bioactive IL-12 and the transpresentation of IL-15/IL15Rα from cDC1 to NK cells. Physical contacts permit delivery of GM-CSF from NK cells to cDC1, which triggers CCR7 on cDC1, instructing them to join the T cell area of the spleen. Abrogation of CCR7 upregulation upon XCR1 genetic inactivation led to a reduced/delayed antiviral CD8+ T cell response, and a failure to control the infection. We also found that, akin to cDC2 (Yi and Cyster, 2013; Gatto et al., 2013), cDC1 upregulate the EBI2/Gpr183 chemokine receptor, which binds 7α,25-OH expressed by stromal cells of the bridging channel (Yi et al., 2012), promoting cDC1 relocalization into the T cell zone. A general schema summarizes our findings (Fig. 8).

**Figure 8:**
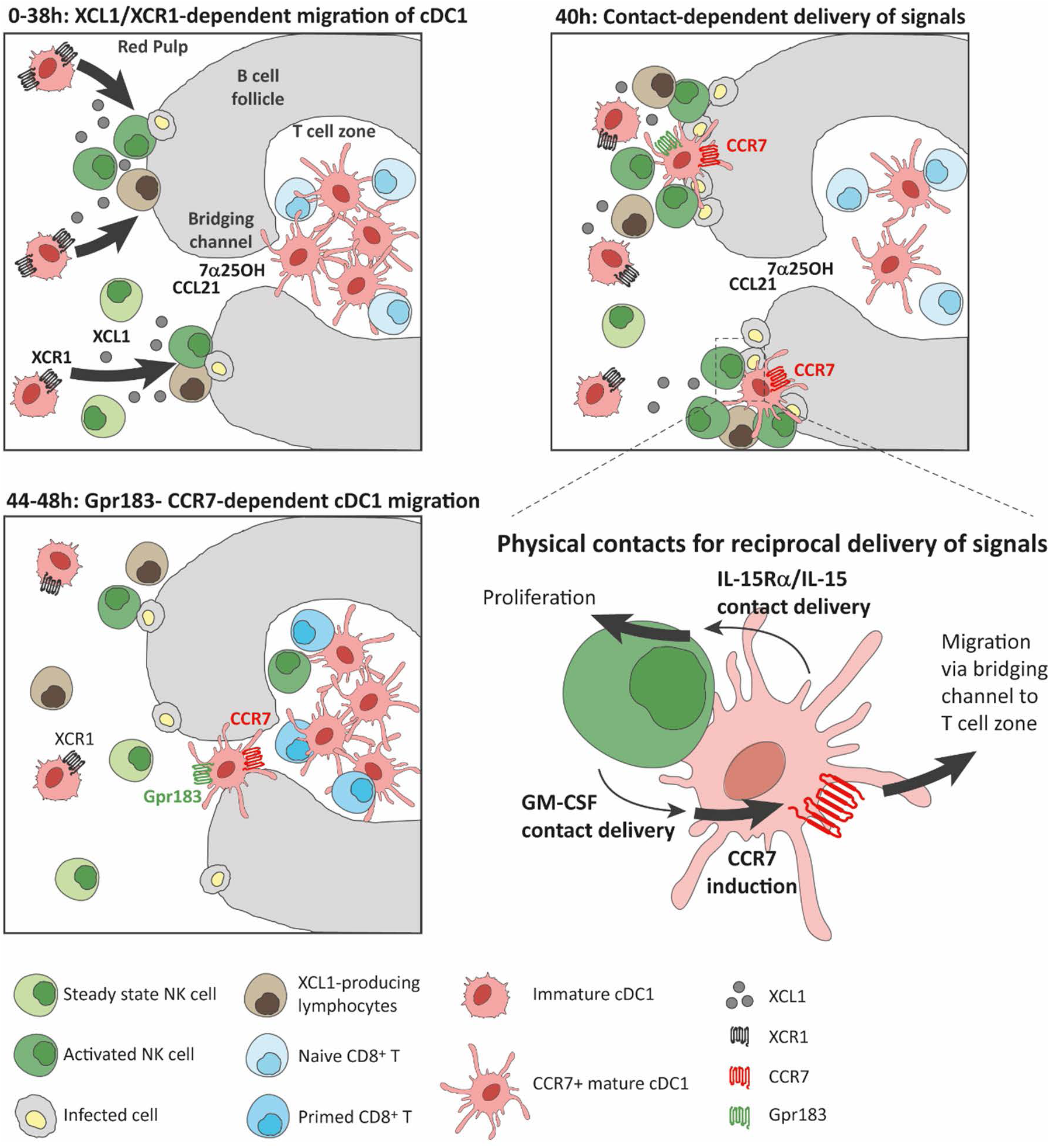
Fine regulation of cDC1 repositioning within the spleen requires physical interactions with NK cells in the marginal zone and reciprocal signal delivery to accelerate the priming of antiviral CD8^+^ T cell responses. Within 38h following viral infection, NK cells and innate-like lymphocytes, positioned near infected cells in the marginal zone, attract red pulp immature cDC1 through XCL1/XCR1 axes. At 40h, the formation of superclusters of activated NK cells and cDC1 in the marginal zone favors physical interactions between these two cell actors, feeding a reciprocal positive autoregulation loop, wherein cDC1 deliver IL-12 and IL-15/IL-15Rα signals to NK cells, promoting their GM-CSF production and expansion respectively. In return, NK cell delivery of GM-CSF to cDC1 triggers CCR7 expression, licensing cDC1 to migrate to the T cell zone. Between 44h and 48h post infection, besides CCR7, cDC1 also acquire the expression of the chemokine receptor Gpr183, which recognizes the bridging channel guiding cue 7α,25-Dihydroxy cholesterol (7α25OH). In T cell zone, CCR7^+^ mature cDC1 prime naïve CD8^+^ T cells, promoting a rapid and efficient adaptive antiviral immune response.

Our study goes beyond the critical role of XCR1 in the spatiotemporal orchestration of cDC1 and NK cell responses during a viral infection. We demonstrate for the first time that NK cells, by providing to cDC1 guiding cues and other signals, finely control their micro-anatomical redistribution within a lymphoid organ, guaranteeing a rapid and protective adaptive antiviral immunity. Because NK cells are the main source of XCL1, we investigated whether they were non-redundant in providing this guiding cue to cDC1. We found that XCL1 selective genetic inactivation in NK cells does not phenocopy XCR1 deficiency (Fig. 3E). This indicates that other cells, such as γδ T, memory T, and NKT cells compensate for the loss of NK cell ability to produce XCL1. Although skin and intestinal γδ T cells are able to upregulate *Xcl1* efficiently upon activation (Boismenu et al., 1996; Rezende et al., 2019), these cells represent only a minute fraction of XCL1-producing cells in the spleen upon MCMV (Fig. 3B and 3D). In contrary, memory αβ T cells and NKT cells account for a significant proportion of the cells expressing XCL1 in the spleen of MCMV-infected mice. Interestingly, it was also observed in other infectious models that activated and memory CD8^+^ T cells and NKT cells could relocalize in cell clusters at the edge of the marginal zone (Alexandre et al., 2016; Bajenoff et al., 2010; Barral et al., 2012), together with NK cells (Alexandre et al., 2016). Therefore, clusterized memory αβ T cells and NKT cells may provide XCL1 and act together to collectively attract cDC1 in their vicinity. Interestingly, XCR1 endows cDC1 with migratory features only during immune responses against viruses, such as MCMV infection (this study) or Vaccinia (Brewitz et al., 2017). In context of cancers, however, XCR1 remained individually dispensable in promoting recruitment of cDC1 for the activation of protective effector lymphocytes, even in tumors that are highly inflammatory (Böttcher et al., 2018), suggesting that the XCL1/XCR1 axis may be functionally conserved to promote a rapid defense in case of acute threats rather than chronic pathologies.

Lymphoid cells, in particular NK cells or ILC3, are known to secrete large amounts of GM-CSF (Dougan et al., 2019). We found that MCMV infection also induces GM-CSF production by NK cells (Fig. 6B) most likely through IL-12 secretion by DCs (Fig. 6C). Members of the IL-12 family cytokines seem to provide a major signal for GM-CSF induction by ILCs, as IL-23 triggers GM-CSF production by ILC3 in colon upon CD40-induced colitis (Pearson et al., 2016). The role of NK cell GM-CSF has been largely overlooked until recently. Louis and colleagues have found that GM-CSF production by NK cells feeds an inflammatory condition by increasing life span and functions of neutrophils and macrophages during autoantibody-mediated inflammatory arthritis (Louis et al., 2020). NK cell GM-CSF was also suggested to promote neutrophil antifungal activity in a model of *Candida albicans* infection (Bär et al., 2014). For a long time, GM-CSF function was mostly investigated with regards to myeloid cell development and differentiation at steady state and upon inflammation (Dougan et al., 2019). At steady state, however, GM-CSF also promotes the differentiation of cDC1 only in peripheral tissues that are constantly under microbiota stimulation such as skin or intestine, but has no major impact on lymphoid tissue-resident cDC1 (King et al., 2010; Greter et al., 2012). Our study goes further in describing a new critical function for GM-CSF on splenic cDC1 in a mildly inflamed environment. We showed for the first time that, probably through cell/cell contacts, NK cell-derived GM-CSF drives CCR7 expression on cDC1, thus instructing their migration in the spleen upon viral infection. Interestingly, GM-CSF does not trigger the expression of the co-stimulatory molecule CD86 on cDC1 in our experimental setting, contrary to what has been reported before in *in vitro* FLT3-L-mediated cDC1 differentiation from BM cell cultures (Mayer et al., 2014). This may be explained by differential DC exposures to GM-CSF concentrations and durations, or differences in cellular microenvironment inherent to in vitro vs in vivo models.

The activated NK cells from MCMV-infected mice do not promote a global maturation program in cDC1, as indicated by the unchanged expression of CD86, but seem to rather enhance CCR7 expression on cDC1 (Fig. 5). Therefore, by regulating cDC1 repositioning toward naïve CD8+ T cells, GM-CSF-producing NK cells indirectly boost antiviral CD8^+^ T cell priming (Fig. 7). This observation contributes to advance our understanding of how, in hosts in whom the early phase of NK cell response is highly effective, activation of antiviral CD8^+^ T cells occurs faster than in hosts with defective NK cells (Robbins et al., 2007). NK cells can functionally affect the magnitude and quality of adaptive immune responses through direct and indirect pathways. NK cell secretion of cytokines, such as IFN-γ or NK cell direct cytolytic activity, may regulate directly T cell responses (Crouse et al., 2015). NK cells also modulate indirectly T cell priming by promoting DC specific functions. Through killing target cells, NK cells provide cDC1 with apoptotic cell-derived antigens, which are cross-presented to naïve CD8^+^ T cells (Krebs et al., 2009; Iyoda et al., 2002; Deauvieau et al., 2015; Dao et al., 2005). NK cells have been shown to induce DC maturation, although the specific molecular mechanisms that came into play remained elusive (Gerosa et al., 2002). Here we unravel how NK cells orchestrate the spatiotemporal repositioning of cDC1 in regions where they have more chance to prime naïve CD8^+^ T cells, an aspect that has never been described before, although the regulation of DC migration by ILCs has been suggested recently (Böttcher et al., 2018).

We find that ILCs, besides attracting cDC1, license them to migrate to the micro-anatomical area in tissue where T cells reside in spleen. This cDC1 relocalization optimizes the relay between innate and adaptive immunity, by accelerating and potentiating T cell expansion and effector functions, therefore yielding a more robust resistance to infections. Whether XCR1/XCL1 and ILC GM-CSF mechanisms are also critical to license tissue-resident cDC1 to migrate into the draining-lymph nodes remains to be determined.

## Material and Methods

### Mice

C57BL/6J and BALB/cByJ mice were purchased from Charles River Laboratories. *Xcr1*^-/-^ (B6.129P2*-Xcr1*^*tm1Dgen*^; MGI:3604538) were backcrossed >10 times into C57BL/6J background before use. C57BL/6J mice or, when possible, littermates were used as controls. *Karma-tdTomato-hDTR* (*Gp141b*^*tm1Ciphe*^) were treated with diphtheria toxin (DT) as previously described (Alexandre et al., 2016). *Karma-Cre* (B6-*Gpr141b*^*tm2Ciphe*^) mice (Mattiuz et al., 2018) were first bred with *Rosa26*^*lox−stop−lox−tdRFP*^ (*Gt(ROSA)26Sor*^*tm1Hjf*^) (Luche et al., 2007) to give *Karma-Cre;Rosa26*^*tdRFP*^, then with *Xcr1*^-/-^ mice to allow the tracking of *Xcr1*-deficient cDC1 in the spleen. *Xcr1*^+/-^ littermate controls were used. For some experiments, *Karma-Cre;Rosa26*^*tdRFP*^ mice were bred to *Il12eYFP*^*+*^ (B6.129*-Il12b*^*tm1*.*1Lky*^/J (Reinhardt et al., 2006) backcrossed >10 times to C57BL*/*6J). To selectively inactivate *Il15ra* gene in cDC1, we followed a specific breeding schema, which prevents germline recombination, using *Xcr1-Cre* (B6-*Xcr1*^*tm1Ciphe*^) (Mattiuz et al., 2018), *Il15ra* full knock-out *Il15ra*^*ko/ko*^ (*Il15ra*^*tm1Ama*^) and *Il15ra*^*flox/flox*^ (*Il15ra*^*tm2*.*1Ama*^) mice (Mortier et al., 2009). We bred *Xcr1*^*cre/cre*^;*Il15Ra*^*ko/ko*^ mice to *Il15ra*^*flox/flox*^ to obtain *Xcr1*^*Cre/wt*^;*Il15Ra*^*fl/ko*^ mice that we experimentally compared to *Il15Ra*^*fl/ko*^ controls. The *Xcl1-mTfp1*^*fl/fl*^ (*Xcl1*^*tm1Ciphe*^) mice were generated according to a standard gene targeting approach in *C57BL/6N*-derived ES cells. A LoxP-exon3-IRES-mTfp1-LoxP cassette was inserted into the 3′-UTR of the *Xcl1* gene (Fig. S3A). *Xcl1-mTfp1*^*fl/fl*^ mice were backcrossed for >three generations with C57BL/6J mice, then with *Nkp46*^*Cre*^ (*Ncr1*^*tm1*.*1(iCre)Viv*^) mice (Narni-Mancinelli et al., 2011) to inactivate the *Xcl1* gene in NK cells. *Csf2rb*^*-/-*^ (B6.129S1*-Csf2rb*^*tm1Cgb*^*/*J (Robb et al., 1995)) were bred to CD45.1^+^ congenic C57BL/6J (B6.SJL-*Ptprc*^*a*^*Pepc*^*b*^/BoyJ) mice to generate *CD45*.*1*^*+*^*Csf2rb*^*-/-*^ mice. BM cells from these mice were used to generate BM chimera mice. All mice were bred under pathogen-free conditions at the Centre d’Immunophénomique de Marseille-Luminy (CIPHE). Seven to 12 weeks-old mice were used for experiments. The animal care and use protocols (APAFIS #1212 and APAFIS #16547) were designed in accordance with national and international laws for laboratory animal welfare and experimentation (EEC Council Directive 2010/63/EU, September 2010), and approved by the Marseille Ethical Committee for Animal Experimentation (registered by the Comité National de Réflexion Ethique sur l’Expérimentation Animale under no. 14).

### MCMV infection and quantification of viral loads

MCMV infections were performed ip with 1×10^5^ plaques forming units (PFU) of salivary gland-extracted MCMV (Smith strain). At this dose, all *C57BL*/6J mice survived the infection whereas all *BALB*/c died between 5 and 6 dpi (not shown). For viral loads, 5 days after infection, total spleens and pieces of livers were harvested. Pieces of livers were weighted before proceeding to conventional plaque assay on organ homogenates as described previously (Crozat et al., 2007).

### Poly(I:C) administration, *in vivo* NK cell depletion and IFN-γ blocking

50ug of the TLR3 analog poly(I:C) (high molecular weight, InVivogen) was administrated iv. For *in vivo* depletion of NK1.1-expressing cells, mice were injected ip the anti-NK1.1 antibody clone PK136 (200ug) or isotype control 12h before poly(IC) administration. To block IFN-γ *in vivo*, 1mg of the anti-IFN-γ clone XMG1.2 was administrated ip to mice 7h prior to infection. Both PK136 and XMG1.2 were produced in house.

### Spleen preparation and flow cytometry analysis

Mice were sacrificed at indicated times post-infection and spleens harvested. For lymphocytes analysis, spleens were smashed through nylon meshes in complete RPMI medium, red blood cells lysed, and filtered again through a nylon mesh before proceeding to antibody stainings. For DC analysis, spleens were digested as previously described (Mattiuz et al., 2018). For the analysis of cytokine induction in DCs, 10µg/ml Brefeldin A was added to the enzymatic cocktail to avoid vesicle release of the cells during the incubation at 37°C. Stained cell acquisition was performed on a FACSCANTO II or a LSR II flow cytometer (BD Biosciences). cDCs were defined as CD3e^-^, NK1.1^-^, CD19^-^, CD11c^high^, SiglecH^-^, and further split into cDC1 and cDC2 which were defined as CD11b^-^CD8α^+^ versus CD11b^+^, CD8α^-^, since XCR1 as a defining marker could not be used in experiments involving *Xcr1*^-/-^ mice. pDCs were defined as CD3e^-^, NK1.1^-^, CD19^-^, CD11c^med^, SiglecH^+^.

### Immunohistofluorescence

Spleens of *Karma*^*Cre*^;*Rosa26*^*tdRFP*^;*Il12eYFP*^*+*^, *Karma*^*Cre*^;*Rosa26*^*tdRFP*^;*Xcr1*^*-/-*^ and *Karma-* ^*Cre*^;*Rosa26*^*tdRFP*^;*Xcr1*^*+/-*^ mice were fixed for at least 2 h in Antigenfix (DiaPath) at 4°C, then washed in phosphate buffer (PB1X: 0.02 M NaH_2_PO_4_ and 0.08 M Na_2_HPO_4_) for 1 h. Finally, they were dehydrated in 30% sucrose overnight at 4°C before being embedded in optimal cutting temperature (OCT) freezing media (Sakura Fineteck). 8-12-µm frozen spleen sections were cut with a Leica CM3050 S cryostat. For the staining, sections were blocked in PB1X containing 0.2% saponin, 1% Protein Block (Abcam), and 2% 2.4G2 supernatant, and stained in PB1X, 0.2% saponin. We used an anti-RFP antibody (Rockland) and a secondary donkey anti-rabbit antibody (Jackson Immuno Research) to amplify the tdRFP signal. When the staining required further blocking steps, normal goat or rabbit sera were added. Stained sections were mounted in ProLong Gold antifade reagent (Invitrogen), and acquired using a spectral mode on a LSM780 confocal microscope (Carl Zeiss).

### Quantification of clusters and analysis of cDC1 distribution

We imaged half spleen sections, and randomly mixed them using the random number generator (www.dcode.fr). Three independent manipulators blindly counted clusters using Zen software with a zoom set between 30 and 40%. One cluster was identified as the gathering of at least 10 IFN-γ^+^ cells including NK cells around at least one MCMV (IE-1^+^) stained cell, in the marginal zone. The cluster quantifications were normalized by the section area (mm^2^). Results showed the mean of the three independent counts. To quantify computationally the presence of cDC1 in these clusters, we isolated each clusters using the Region of Interest (ROI) tools on Zen software. Then all ROI were exported as unmodified uncompressed TIFF files and tdRFP intensity was quantified in each ROI using ImageJ, and normalized to the area of the ROI (mm^2^). To follow cDC1 relocalization in the marginal zone, bridging channel and T cell area, we employed a high throughput acquisition of whole spleen sections using the Pannoramic Scan slide scanner (3dHistec). We quantified tdRFP intensity minus background in each of the three areas we identified using MOMA-1 and B220 staining as shown in Supplementary Fig.5. Between 6 to 14 white pulp follicles were analyzed per mouse. The 3-dimensional (3D) reconstruction and the video S1 were generated using the Imaris software (Oxford Instruments).

### Co-culture experiments and migration assays

For co-culture experiments, splenic cDC1 and NK cells were enriched using Miltenyi Biotec CD8^+^ Dendritic Cell isolation kit and NK cell isolation kit respectively. NK cells were isolated from non-infected and 40h-infected mice. For co-culture using bone marrow-derived eqcDC1, cultures of Wt and *Csf2rb*^*-/-*^ bone marrow cells with FLT3-L were performed as described previously (Mattiuz et al., 2018; Hochrein et al., 2004). 14 days later, cells were harvested and enriched for eqcDC1 by removing magnetically SIRPα-expressing cells using the Dynabeads system (ThermoFisher). The enriched cell populations were co-cultured for 6h at a ratio of 1DC:3NK, stained and live cells were analyzed by flow cytometry. We used 0.4um pore Corning Transwells to evaluate the contribution of cDC1/NK physical contacts to CCR7 upregulation. For migration assay, splenic DCs were enriched by OptiPrep (Sigma-Aldrich) and migration experiments were conducted as described previously (Crozat et al., 2010) using 10nM of the 7α,2-hydroxycholesterol (7α,25-OH) (Sigma-Aldrich).

### Microarray experiments and analysis

Each cell type studied was FACS sorted from spleen to over 98% purity in independent duplicate samples. Microarray (MoGene 1.0 ST) experiments were performed as previously published (Robbins et al., 2008), and data were analyzed using the affy and oligo packages in R (version 3.2.2) as previously described (Baranek et al., 2012). Differentially expressed genes were selected using linear models from Limma package in Bioconductor. The microarray data have been deposited in the GEO database under the series accession number GSE142402.

### Statistical analyses

Statistical analyses were performed using a nonparametric Mann-Whitney test performed with Prism 6 (GraphPad Software) for all experiments unless specified otherwise. *n*.*s*., non-significant (p > 0.05); *p < 0.05; **p < 0.01; ***p < 0.001.

## Acknowledgements

We acknowledge France Bio lmaging infrastructure supported by the Agence Nationale de la Recherche (ANR-10-INBS-04-01, call “Investissement d’Avenir”), and in particular Sébastien Mailfert, Roxanne Fabre, Mathieu Fallet and Lionel Chasson from the CIML Immagimm and histology core facilities for technical help. The authors thank Eric Vivier (CIML) for providing *Nkp46-Cre* mice, Sandrine Sarrazin and Michael Sieweke (CIML) for *Csfr2b*-deficient mice, and Mauro Gaya (CIML) for providing CD1d tetramer. This work benefited from institutional supports from CNRS and Inserm, and was funded by the I2HD CIML-SANOFI project, by grants from Fondation pour la Recherche Médicale (label “Equipe FRM 2011”, #DEQ20110421284 to MD) and from the European Research Council under the European Community’s Seventh Framework Programme (FP7/2007-2013 grant agreement 281225, to M.D.), by the XCR1DirectingCells ANR Grant (KC), and by the Fondation ARC pour la Recherche sur le Cancer (KC). SG was supported by a doctoral fellowship from the French Ministère de l’Enseignement Supérieur et de la Recherche, and from Fondation ARC pour la Recherche sur le Cancer. JCC was supported by fellowships from La Ligue Nationale Contre le Cancer.

## Author contributions

SG designed and performed experiments, analyzed, interpreted data, and contributed to manuscript preparation; MA designed, performed, quantified and analyzed data related to microscopy imaging, and contributed to manuscript preparation; JCC performed experiments and analyzed data; MM analyzed microarray data; HL provided critical technical support for imaging; MD contributed to direct and fund the project, and edited the manuscript. KC contributed to conceptualization, directed and contributed to fund the project, designed and performed experiments, analyzed, interpreted data, and wrote the manuscript.

## Conflict of interest

The authors declare no conflict of interests.

## Notes

### Competing Interest Statement

The authors have declared no competing interest.

