## Supplementary material for "NK cells orchestrate splenic cDC1 migration to potentiate antiviral protective CD8+ T cell responses"

Supplemental figure S1

(Related to Fig. 1)

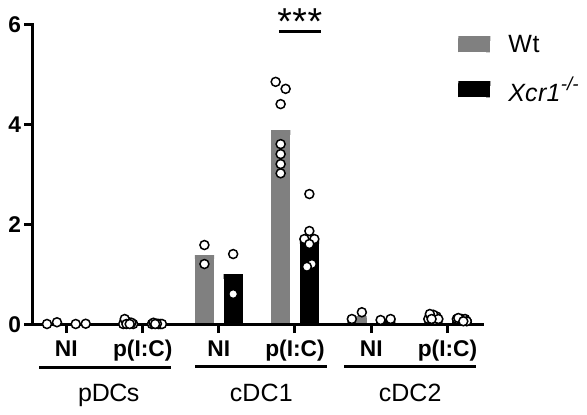

A.

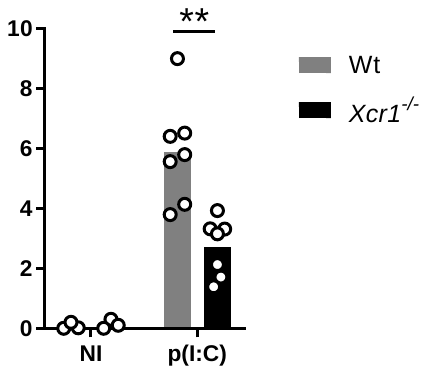

C.

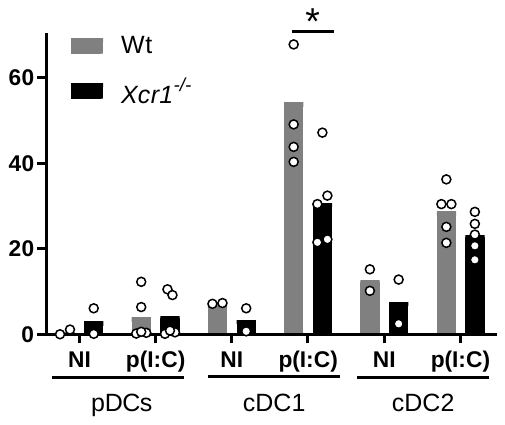

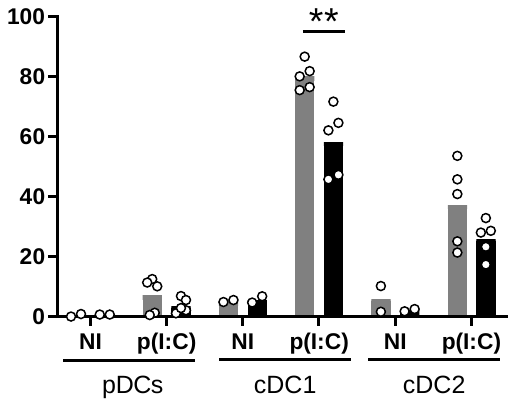

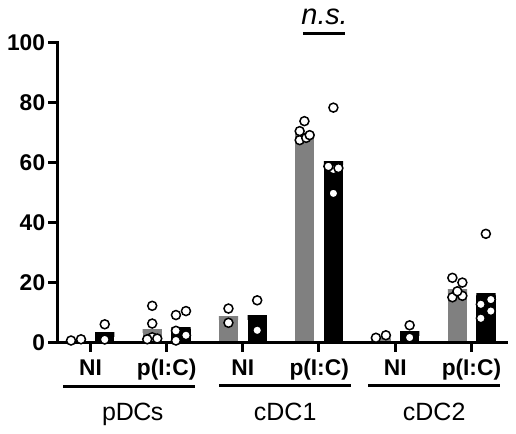

E.

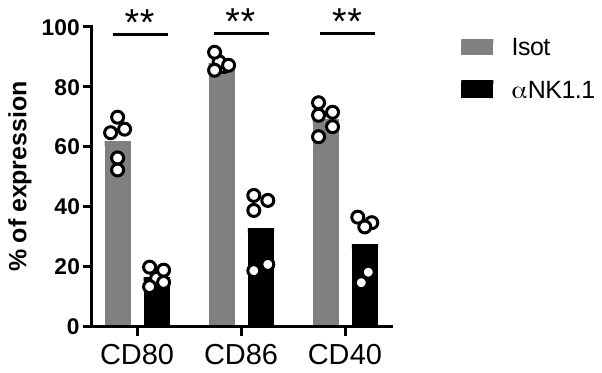

B.

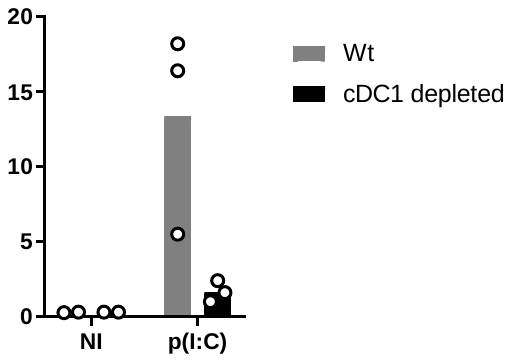

D.

**% IFN-γ+**

**% IL-12p40**

**% IFN-γ+**

**% CD80+**

**% CD86+**

**% CD40+**

NKp46

CD169/MOMA-1

F.

*Xcr1^+/-^*

100 μm

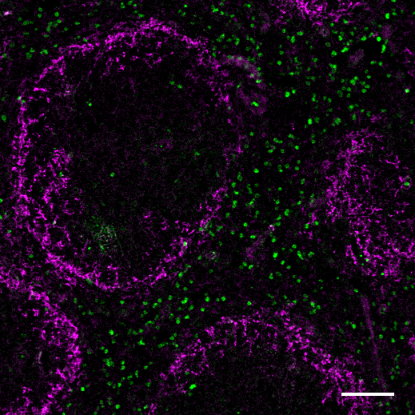

TCZ

WP

RP

MZ

100 μm

BC

**Figure S1 (related to Fig.1): XCR1 promotes mutual activation of cDC1 and NK cells upon poly(I:C) administration. A)** IFN-γ production by NK cells in cDC1-depleted mice 3h post-poly(I:C) injection. *Karma-tdTomato-hDTR* (cDC1 depleted) and Wt controls were injected with 32ng/g of body weight of DT 24h prior poly(I:C) administration. One experiment representative of two independent ones with 3 mice per injected group is shown. NI, non-injected. **B)** cDC1 maturation in the spleen of NK1.1^+^ cell-depleted animals 20h post poly(I:C) administration. Two independent experiments with 2-3 mice per group were pooled. Isot, Isotype; αNK1.1, anti-NK1.1; **, p<0.01. **C-D)** IL-12p40 induction in pDCs, cDC1 and cDC2 **(C)** and IFN-γ production by NK cells **(D)** in Wt vs *Xcr1*-deficient spleens 3h post-poly(I:C) administration. Splenic DCs (CD19^-^TCRβ^-^NK1.1^-^CD11c^lo/hi^) were gated to define pDCs (CD11c^lo^SiglecH^+^), cDC1 (CD11c^hi^SiglecH^-^CD8a^+^CD11b^-^) and cDC2 (CD11c^hi^SiglecH^-^CD8a^-^CD11b^+^). Two independent experiments with at least 3 mice per injected group were pooled. NI, non-injected. ***, p<0.001; **, p<0.01. **E)** Analysis of DC maturation in Wt vs *Xcr1*-deficient spleens 20h post poly(I:C) administration. Two independent experiments with three mice per injected group were pooled. NI, non-injected. *, p<0.05; **, p<0.01, *n.s.*, non-significant. F) Spleen sections of steady state *Karma^Cre^;Rosa26^tdRFP^* were stained for MOMA-1/CD169 (purple) and NKp46 (green). RP, red pulp; MZ, marginal zone; WP, white pulp; BC, bridging channel; TCZ, T cell zone. Scale bar, 100μm. One image representative of 12 analyzed mice from 6 independent experiments is shown.

**Video S1: 3D reconstruction of Fig. 2F micrograph using Imaris.** Color code: IL-12eYFP, yellow; Nkp46, green; IFN-γ, purple; tdRFP, red. Scale bar, 10μm.

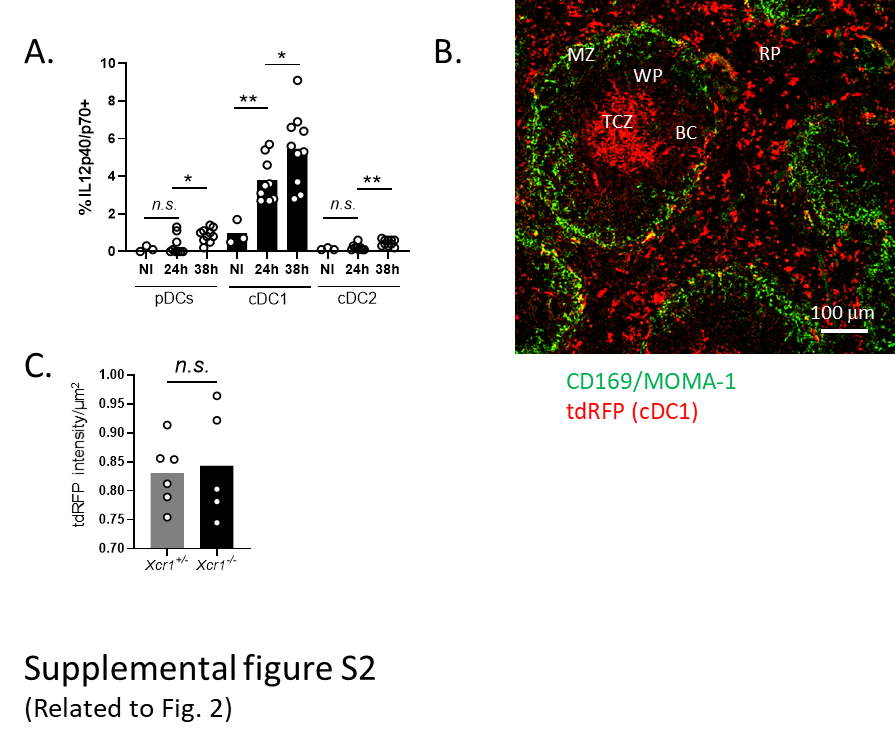

**Figure S2 (related to Fig.2): Analysis of IL-12 production by cDC1 upon MCMV and of cDC1 localization in the spleen at steady state. A**) Spleens were harvested at 24 and 38h post MCMV infection to analyze IL-12p40/p70 production by DCs in Wt animals. Three independent experiments with at least 3 mice per infected group were pooled. NI, non-infected; *n.s*., non-significant; *, p<0.05; **, p<0.01**.** B) Analysis of cDC1 distribution in the spleen at steady state. *Karma^Cre^;Rosa26^tdRFP^* spleen sections were stained for MOMA-1/CD169 (green) and tdRFP (red). RP, red pulp; MZ, marginal zone; WP, white pulp; BC, bridging channel; TCZ, T cell zone. One image representative of 12 analyzed mice from 6 independent experiments is shown. C) Mean fluorescent intensity of the tdRFP per μm^2^ quantified on whole spleen sections analyzed in Fig. 3D. Two independent experiments with 2-3 mice per infected group were pooled. *n.s*., non-significant.

STOP

codon

LoxP

LoxP

Ires

EX 1

EX 2

EX 3

START

codon

5’ UTR

3’ UTR

mTfp1

STOP

codon

EX 1

EX 2

EX 3

START

codon

5’ UTR

3’ UTR

LoxP

LoxP

Ires

STOP

codon

EX 3

mTfp1

***Wt* locus**

**Construct**

***Xcl1-mTfp1^fl/fl^***

**locus**

5’ UTR

3’ UTR

Supplemental figure S3

(Related to Fig. 3)

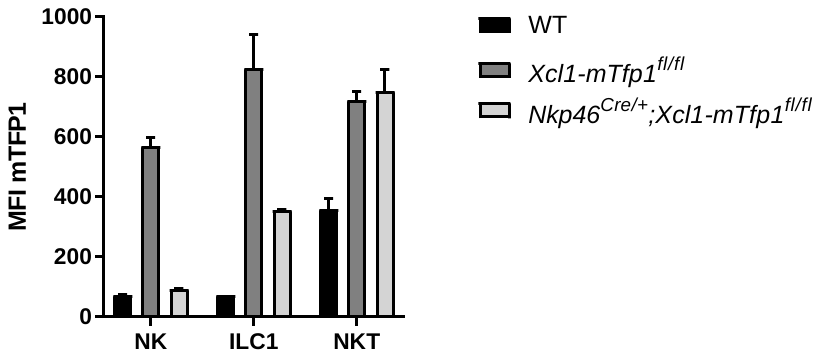

A.

B.

**Figure S3 (related to Fig. 3): Construction of *Xcl1-mTfp1^fl/fl^* mice and analysis of *Xcl1* inactivation in ILCs.** A) The *Xcl1-mTfp1^fl/fl^* mouse model was generated by inserting a LoxP-exon3-IRES-mTfp1-LoxP cassette in frame of the *Xcl1* gene. The *mTfp1* gene codes for the monomeric Teal fluorescent protein mTFP1 (Ex:462; Em:492). In this mouse, the expression of the *Xcl1* locus resulted in an *Xcl1-Ires-mTfp1* transcript, which led to the production of two distinct proteins: XCL1 and mTFP1. B) Analysis of the geometric mean of the mTFP1 fluorescence in ILCs of Wt, *Xcl1-mTfp1^fl/fl^* and *Nkp46^Cre/+^;Xcl1-mTfp1^fl/fl^* mice. One experiment representative of two independent ones with at least 2 mice per group is shown. Error bars represent standard deviations.

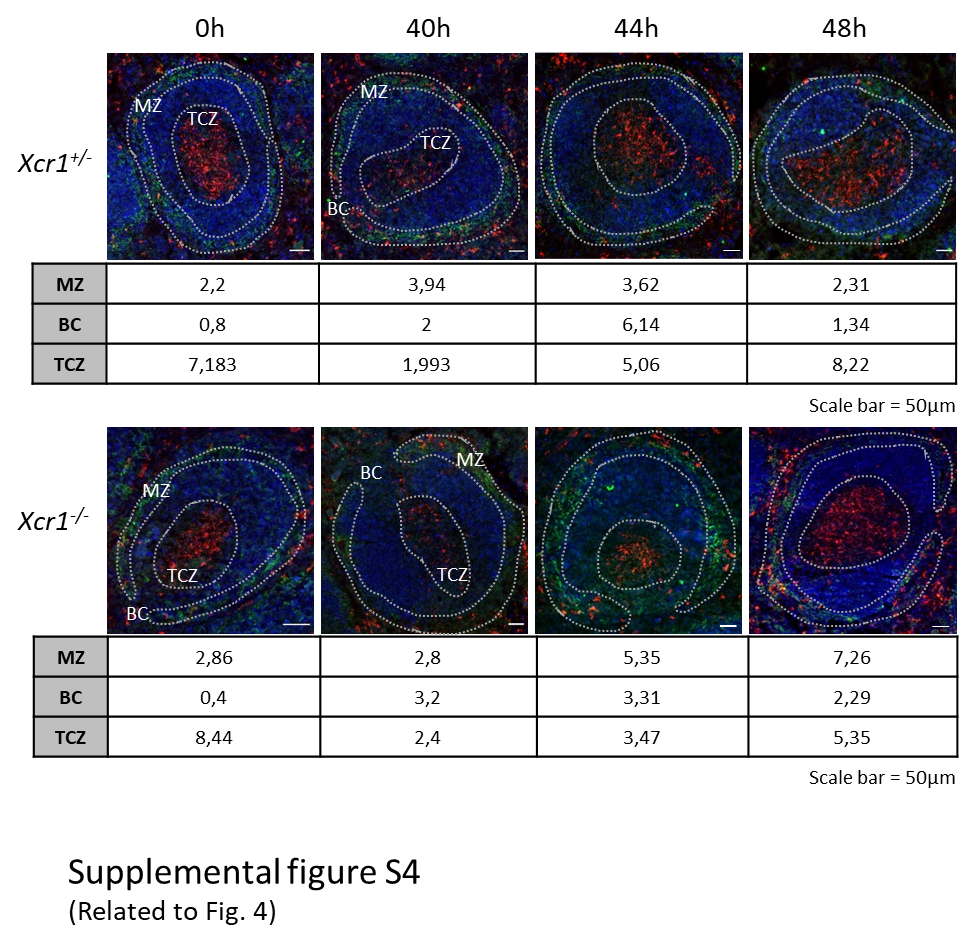

**Figure S4 (related to Fig.4): tdRFP quantification in different regions of the spleen during MCMV infection.** Whole spleen sections of *Karma^Cre^;Rosa26^tdRFP^;Xcr1^-/^****^-^*** mice and *Karma^Cre^;Rosa26^tdRFP^;Xcr1*^+/-^ controls were prepared at indicated time after MCMV infection, and stained for MOMA-1/CD169 (green), B220 (dark blue), tdRFP (red) and nuclei (not shown), before being scanned with Pannoramic Scan slide scanner. The tdRFP intensity per pixel^2^ quantified in each sample shown here has been reported in a table below each respective image. The quantification method has been detailed in the material and methods. MZ, marginal zone; TCZ, T cell zone; BC, bridging channel.

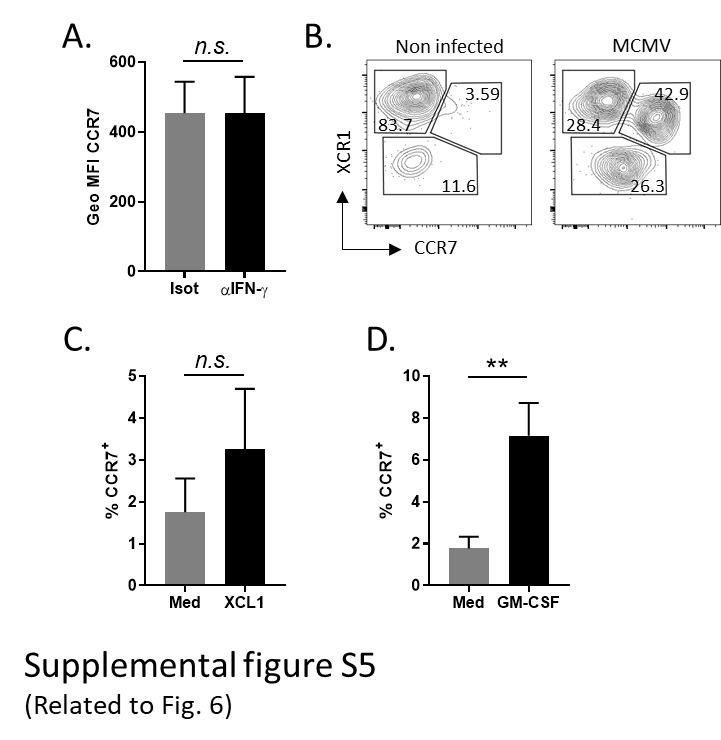

**Figure S5 (related to Fig. 6): Effects of IFN-γ, XCL1, and GM-CSF on CCR7 upregulation on cDC1.** A) CCR7 expression on cDC1 after IFN-γ *in vivo* blocking, 48h post MCMV. One experiment representative of two independent ones with 4 mice per group is shown. *n.s.*, non-significant. B) CCR7 and XCR1 expression analysis on splenic cDC1 48h after MCMV infection. One experiment representative of 3 independent ones with at least 3 mice per infected group is shown. C) CCR7 expression on eqcDC1 differentiated from FLT3-L-cultured bone marrow cells 6h after treatment with recombinant XCL1. Two experiments each with 2-3 mice per group were pooled. *n.s.*, non-significant. D) Analysis of CCR7 induction on eqcDC1 differentiated from FLT3-L-cultured bone marrow cells 6h after treatment with recombinant GM-CSF. Three experiments each with 3-4 mice per group were pooled. **, p<0.01. Error bars represent standard deviations.

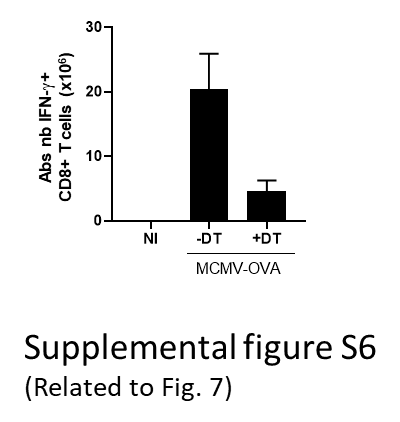

**Figure S6 (related to Fig.7): cDC1 are critical for the expansion of virus-specific CD8^+^ T cells.** Analysis of OVA-specific CD8^+^ T cell expansion in cDC1-depleted (+DT) *vs* non-depleted (-DT) mice 5 days post-MCMV-OVA infection. DT was administrated to *Karma-tdTomato-hDTR* mice 1d prior to infection, and 2.5 d after. Splenocytes were incubated for 4h with SIINFEKL peptide before intracellular IFN-γ staining. One representative experiment of two independent ones with at least 3 mice per group is shown. NI, non-infected.
